# Protein Scaffolds Modulate the Electronic Structure of a Common Ketyl Radical Intermediate in Thiamine Pyrophosphate-Dependent Radical Biocatalysis: Insights from EPR and ENDOR Spectroscopy

**DOI:** 10.64898/2026.10.15.759499

**Authors:** Guodong Rao, Zheng Zhang, Xin Liu, Angela A. Shiau, R. David Britt, Yang Yang

## Abstract

New-to-nature radical biocatalysis has emerged as a powerful strategy for asymmetric C–C bond formation, yet the enzyme-bound radical intermediates that underpin these transformations have remained largely uncharacterized. Here, we combine photobiocatalytic reaction development with multifrequency electron paramagnetic resonance (EPR), electron-nuclear double resonance (ENDOR) spectroscopy, and density functional theory (DFT) calculations to elucidate the ketyl radical intermediates formed during thiamine pyrophosphate (TPP)-dependent radical biocatalysis. We first develop an oxidative photobiocatalytic cross-coupling of aldehydes with alkyltrifluoroborates catalyzed by engineered benzoylformate decarboxylase (BFD) variants, providing enantioenriched α-chiral ketones through a new oxidative radical manifold inaccessible to benzaldehyde lyases (BALs). Freeze-quench multifrequency EPR and ENDOR spectroscopy directly identify the catalytically generated TPP-bound ketyl radical and reveal that this common radical intermediate adopts distinct electronic structures in BFD and BAL despite their identical chemical composition. Analysis of ^1^H, ^19^F, ^31^P, and ^14^N hyperfine interactions, together with DFT calculations, demonstrates that enzyme-specific active-site conformations redistribute spin density between the TPP cofactor and substrate fragment. Greater substrate-centered spin delocalization in BFD provides a mechanistic basis for its superior radical C–C coupling activity relative to BAL. These results establish a direct relationship between protein-controlled electronic structure and catalytic function in TPP-dependent radical biocatalysis and demonstrate that engineering the electronic properties of a common radical intermediate represents a useful strategy for controlling enzyme reactivity in radical biocatalysis.

**Table of Content:** 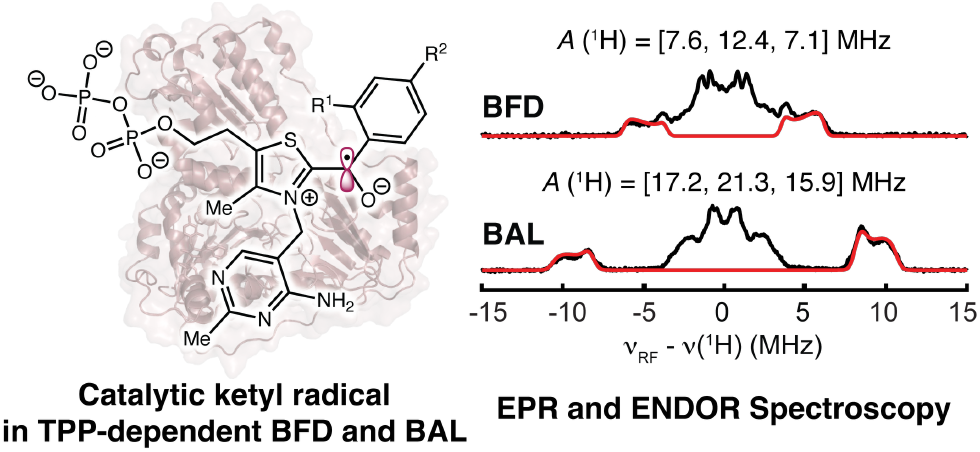

## INTRODUCTION

Over the past decade, non-native radical biocatalysis has emerged as a powerful tool for achieving challenging stereoselective free radical reactions.^1–6^ By leveraging the innate redox properties of various organic^7,8^ and metallocofactors^6,9–13^ or by using a secondary small-molecule catalyst to trigger redox events,^4–6,14–16^ a wide range of co-factor-dependent^17–19^ and cofactor-independent enzymes^20^ has been reprogrammed to catalyze radical reactions through non-native mechanisms. In particular, cooperative catalysis with photoredox catalysts,^21^ operating through enzymatic cofactor-derived covalent intermediates, has enabled organic cofactor-dependent enzymes, including pyridoxal 5’-phosphate (PLP)^18^ and thiamine pyrophosphate (TPP)^19a–i^-dependent enzymes, to be repurposed and engineered to catalyze a wide range of synthetically valuable, asymmetric intermolecular radical reactions. These transformations provide access to a broad spectrum of valuable molecules, including many previously inaccessible through traditional chemical or biological means.

Although biocatalytic free radical transformations using cofactor-dependent enzymes have advanced rapidly, the catalytic mechanisms of the underlying enzymology remain largely hypothetical and lack full experimental validation. In previous mechanistic proposals, radical addition^18,20c,d^ or electron transfer^6,19,20a,b^ to closed-shell enzymatic covalent intermediates was invoked to generate enzyme-bound open-shell covalent intermediates (Scheme 1A), which are central to advancing non-native radical biocatalysis in these enzyme families. However, the transient, highly reactive nature of these enzyme-bound radical intermediates has made their unambiguous characterization by electron paramagnetic resonance (EPR) spectroscopy challenging. Several difficulties complicate the study of these enzymatic radical intermediates. First, the interweaving dual catalytic cycles of cooperative photobiocatalysis, together with the involvement of photochemistry, make it nontrivial to accumulate sufficient quantities of reactive enzymatic radical intermediates for spectroscopic characterization. Second, unlike naturally occurring radical enzymes optimized by natural evolution, most non-native radical enzymes are engineered through mechanism-guided repurposing after only a handful of rounds of directed evolution. As a result, these reprogrammed enzymes have not undergone the extensive adaptive evolution needed to optimize their open-shell reactivity. Consequently, these systems often exhibit slow radical generation kinetics and limited radical stability, which impede full spectroscopic elucidation of these species.

**Scheme 1.**
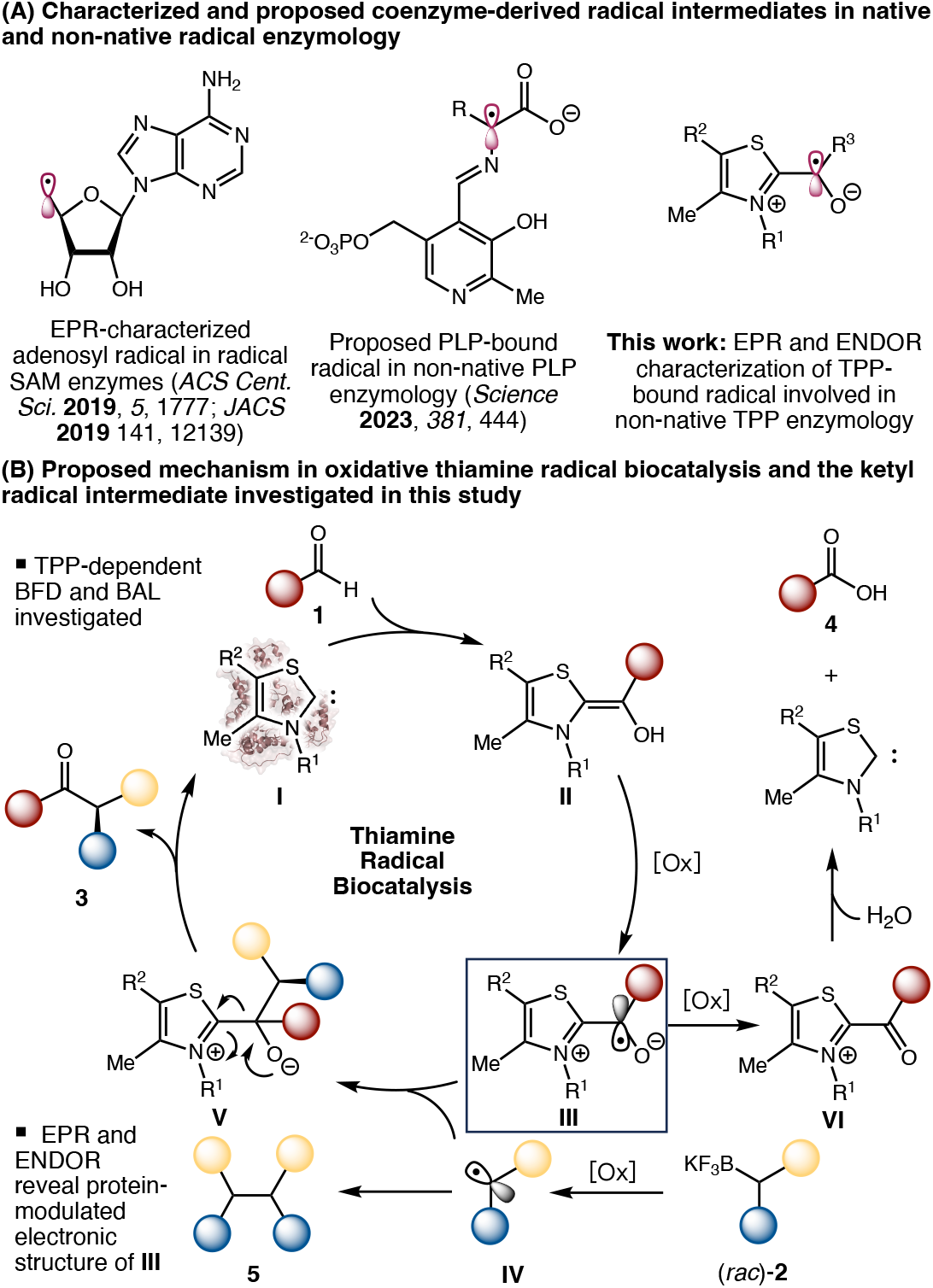
Overview of Coenzyme-Derived Radical Intermediates in Enzymology, Proposed Mechanism in Oxidative Thiamine Radical Biocatalysis and the Ketyl Radical Intermediate Investigated in This Study.

Our UC Davis group and other leading EPR labs have extensively applied EPR spectroscopy to advance the field of radical enzymology, providing fundamental mechanistic insights into naturally occurring radical enzymes, including the *S*-adenosyl-L-methionine (SAM) enzyme superfamily (Scheme 1A).^22–25^ Building on recently developed non-native radical enzymology, we set out to investigate the mechanism and nature of the radical intermediates involved in radical biocatalytic reactions catalyzed by repurposed natural enzymes. We were particularly interested in recently reported radical reactions catalyzed by TPP-dependent enzymes (Scheme 1B).^26^ In 2024, inspired by the pioneering studies by Studer, Rovis, Chi, Scheidt, Ye, and other researchers on small-molecule *N*-heterocyclic carbene (NHC) radical catalysis,^27^ particularly the elegant work of Ohmiya and coworkers,^28,29^ the Huang laboratory^19a^ and our UCSB laboratory^19b^ leveraged cooperative photobiocatalysis to develop the first examples of TPP enzyme-catalyzed asymmetric radical alkylation for the enantioselective synthesis of α-chiral ketones. In these studies, the Huang group employed an engineered benzaldehyde lyase (BAL)^19a^ to convert aldehyde substrates, while our group evolved a benzoylformate decarboxylase (BFD)^19b^ to transform both aldehydes and α-keto acids. Mechanistically, the deprotonated thiazolidine (**I**) reacts with the benzaldehyde substrate to form the Breslow intermediate (**II**). Subsequent single-electron oxidation, either by an excited-state photosensitizer or an appropriate chemical oxidant, converts this Breslow intermediate into a distinct ketyl radical (**III**), which enables radical-mediated C–C coupling and drives product formation through collapse of the tetrahedral intermediate. This critical ketyl radical derived from acetyl-CoA has previously been characterized by EPR and ENDOR spectroscopy in the native radical TPP enzyme pyruvate: ferredoxin oxidoreductase (PFOR), providing mechanistic insights into canonical TPP radical enzymology.^30^ However, the corresponding ketyl radical has not been investigated by modern multifrequency EPR and ENDOR techniques in repurposed TPP-dependent enzymes, whose native function do not involve open-shell intermediates. As a result, the electronic structure and catalytic significance of this ketyl radical species underlying newly developed TPP radical biocatalysis have remained largely underexplored.

Recent elegant studies from Huang,^19a,c–k^ Fasan^19l^ and Hayashi/Kato^19m^ further generalized thiamine radical biocatalysis, leading to a plethora of synthetically useful methods for asymmetric C–C bond formation. In particular, BAL^19a,c,e,f,g,j,k,l^ and BFD^19b,h,i^ have displayed complementary reactivities and remain essential workhorse enzymes in recently developed biocatalytic radical reactions. Despite this progress, the structural and mechanistic basis for the divergent behavior of BAL and BFD—and the factors underlying their contrasting yet complementary reactivities in radical biocatalysis— remain to be elucidated. Herein, we integrate biocatalytic reaction development, mechanistic studies, and EPR spectroscopy to elucidate the nature of enzymatic TPP-bound ketyl radical intermediates and to shed light on the electronic and reactivity differences between the BAL-and BFD-derived ketyl radical species.

## RESULTS AND DISCUSSION

To further generalize TPP-catalyzed asymmetric radical alkylation, we recently investigated the use of easily available organoboron substrates as radical precursors in biocatalytic C–C coupling to prepare enantioenriched α-chiral ketones (Table 1). Compared to the redox-active ester-based C–C coupling process previously reported by the Huang lab and our lab,^19a,b^ this Suzuki-type cross-coupling of alkyltrifluoroborates is an oxidative process and requires the use of a stoichiometric amount of judiciously selected oxidant.

**Table 1.**
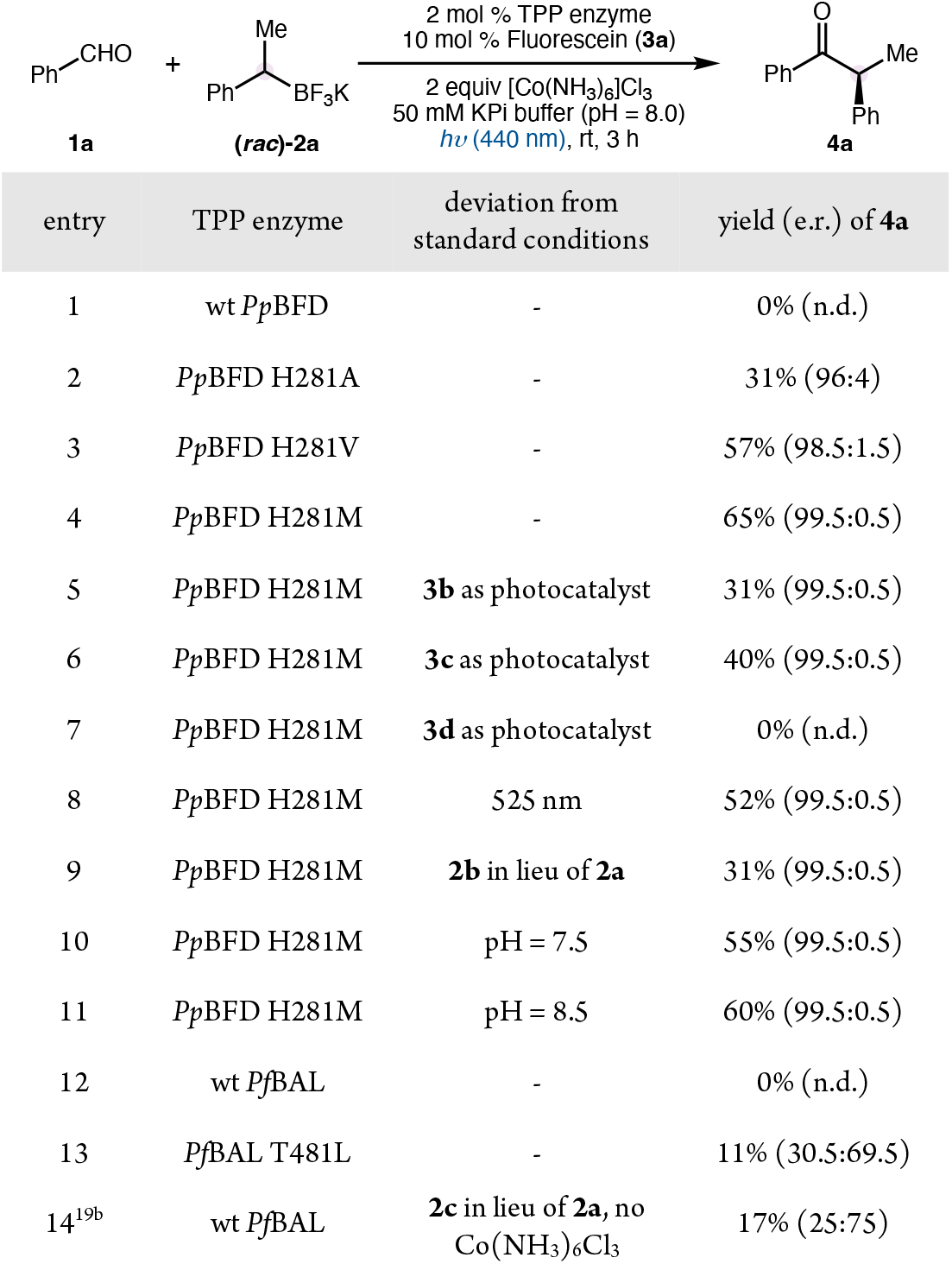
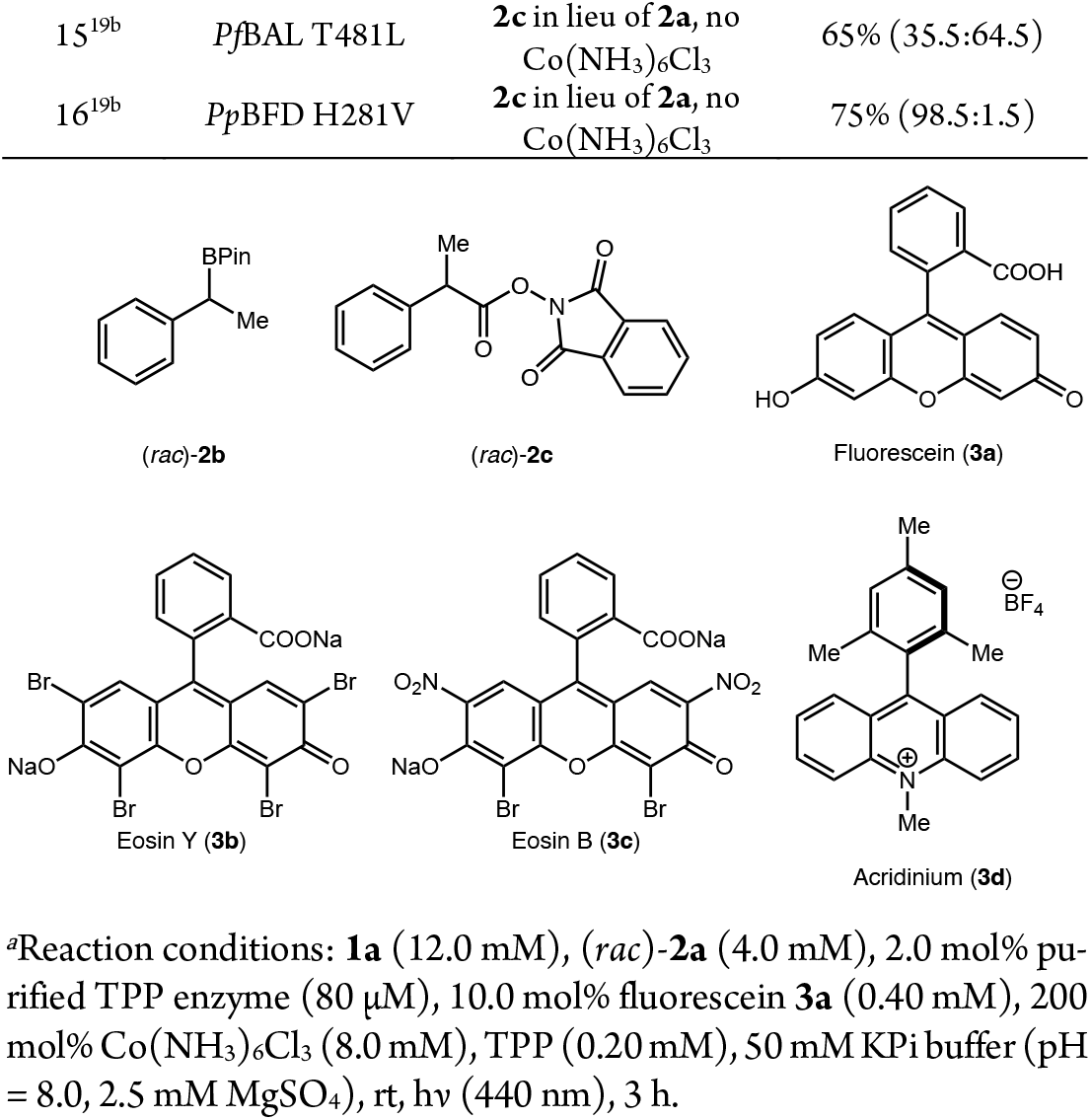
Contrasting TPP Enzyme Effects in Oxidative and Redox-Neutral Radical C–C Coupling.^*a*^.

Using benzaldehyde (**1a**) and racemic secondary alkyltrifluoroborate ((*rac*)-**2a**) as the model substrates, Co(NH_3_)_6_Cl_3_ as the oxidant and fluorescein (**3a**) as the photosensitizer, we evaluated the use of TPP-dependent enzymes previously shown to be effective in the redox-neutral radical coupling. Although wild-type *Pseudomonas putida* BFD (*Pp*BFD) did not provide any desired product **4a** (Table 1, entry 1), the use of its single mutant *Pp*BFD H281A with a broadened substrate entrance tunnel allowed **4a** to form in 31% yield and 96:4 e.r. (entry 2). A survey of other H281 mutants revealed both *Pp*BFD H281V and *Pp*BFD H281M to be more effective mutants, giving rise to **4a** in 57% yield and 98.5:1.5 e.r. (entry 3) and 65% yield and 99.5:0.5 e.r. (entry 4), respectively. Other xanthene-derived photocatalysts, including eosin Y (entry 5) and eosin B (entry 6), led to reduced catalytic efficiency and identical enantioselectivity. Additionally, an acridinium photocatalyst (entry 7) was found to be ineffective in promoting this transformation. Using *Pp*BFD H281M as the biocatalyst and fluorescein as the photocatalyst, we further examined the reaction conditions. Irradiation using a green LED at 525 nm resulted in a slightly lower yield of 52% (entry 8). Replacing the organotrifluoroborate salt **2a** with the corresponding pinacol boronic ester **2b** also provided an inferior yield of 31% (entry 9). Reducing or increasing the buffer pH to 7.5 or 8.5 lowered the yield of **4a** to 55% (entry 10) and 60% (entry 11), respectively. Collectively, these results established the optimal conditions for this photobiocatalytic oxidative radical coupling.

Surprisingly, when wt *Pseudomonas fluorescens* BAL (*Pf*BAL) and *Pf*BAL T481L were used as biocatalysts, no product **4a** (entry 12) or only 11% of **4a** (entry 13) was observed, respectively. The drastically reduced catalytic efficiency of wt *Pf*BAL and *Pf*BAL T481L stood in stark contrast to their demonstrated ability to facilitate the redox-neutral photobiocatalytic coupling of redox-active esters (**2c**). Under those conditions, wt *Pf*BAL and *Pf*BAL T481L furnished **4a** in 17% yield (25:75 e.r.) and 65% yield (35.5:64.5 e.r.), respectively (entries 14 and 15). The contrasting behavior of *Pp*BFD and *Pf*BAL highlighted substantial, previously underappreciated differences in the activity of these TPP-dependent enzymes.

To gain further understanding of the distinct properties of TPP-dependent *Pp*BFD and *Pf*BAL variants in radical catalysis, we employed multifrequency continuous-wave (CW) and pulsed electron paramagnetic resonance (EPR) spectroscopy to characterize the enzyme-bound radical intermediates formed during catalysis. To trap these transient radical species, biocatalytic reactions were performed under turnover conditions containing 1mM TPP-dependent enzymes (BFD or BAL), followed by rapid freeze-quenching for EPR analysis. As shown in Figure 1A, X-band CW EPR spectra revealed the accumulation of radical species in reactions catalyzed by *Pp*BFD H281V and *Pf*BAL T481L. Spin quantification indicated that approximately 30 μM radical accumulated (∼3% of total enzyme concentration) under these conditions. Although both enzymes generated comparable levels of radical intermediates, their EPR lineshapes differ markedly, suggesting distinct electronic structures and/or active-site environments surrounding the enzyme-bound radicals.

**Figure 1.**
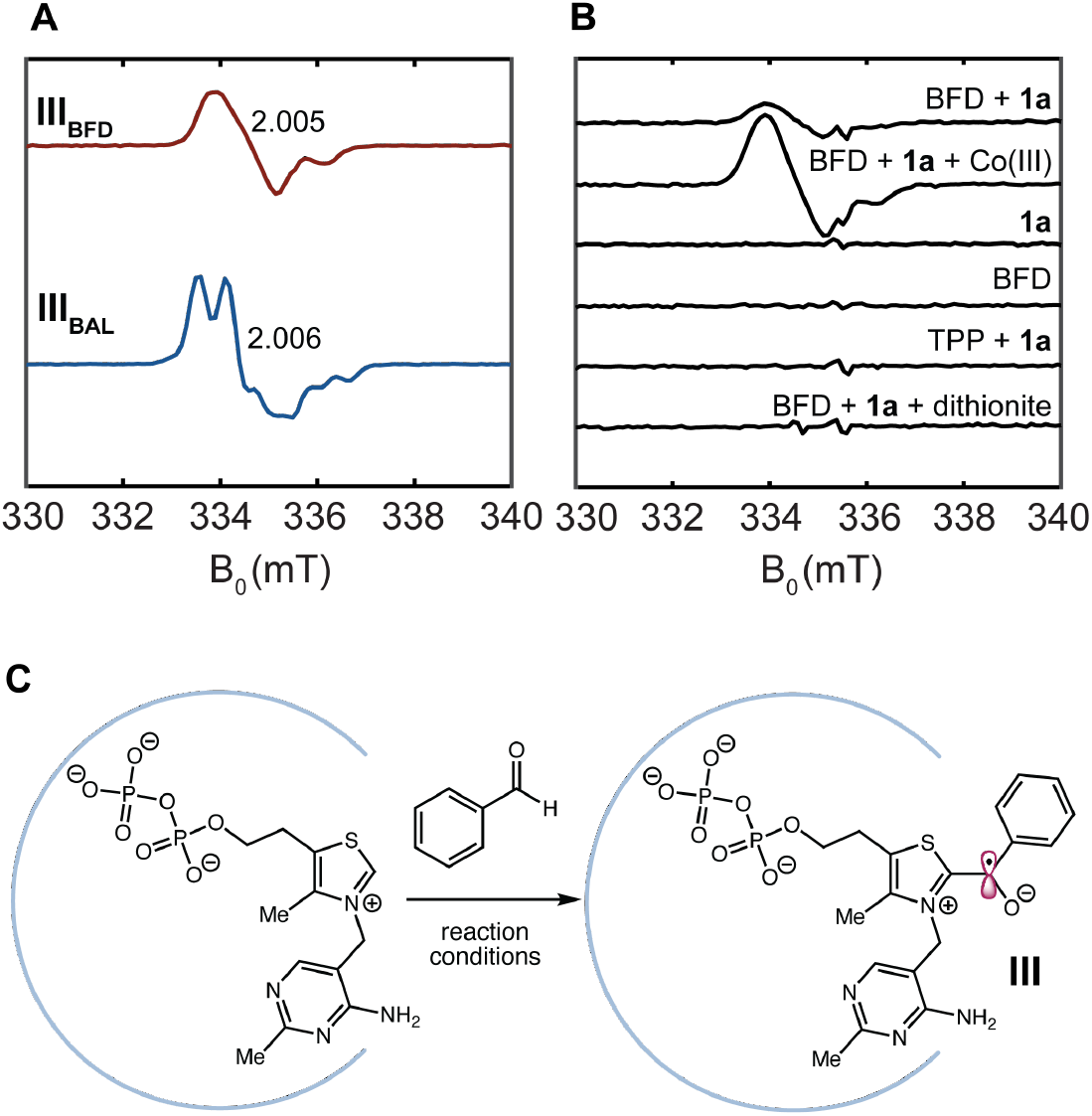
X-band EPR (100 K, 0.2 mW) spectra of radical species generated in the biocatalysis using BFD and BAL. (A) X-band CW EPR spectra of **III**_**BFD**_ and **III**_**BAL**_. (B) Formation of **III**_**BFD**_ under different conditions. (C) Scheme showing the structure and formation of **III**.

To probe the origin of this radical, a series of control experiments was performed (Figure 1B). In both *Pp*BFD H281V and *Pf*BAL T481V, simply mixing the enzyme with substrate **1a** produced the same radical signal, whereas no EPR signal appeared from the enzyme alone, the substrate alone, or the substrate combined with the TPP cofactor alone. These observations indicate that radical formation requires the TPP enzyme active site and is associated with the formation of the TPP-bound Breslow intermediate (**II**). Furthermore, radical accumulation was enhanced by ∼3.5-fold in the presence of Co(III) and completely abolished by sodium dithionite. The small amount of radical observed in the absence of an added oxidant is likely due to spontaneous oxidation of the Breslow intermediate in the presence of trace amount of O_2_, consistent with its strongly reducing nature (*E*^0^ = –0.8 V *vs*. Fc/Fc^+^).^31^ Together, these results support the assignment of the observed species as the one-electron oxidized Breslow intermediate, namely the ketyl radical **III** (Figure 1C, Scheme 1), hereafter referred to as **III**_**BFD**_ and **III**_**BAL**_, respectively.

To assess whether the formation of this enzymatic radical intermediate is substrate-dependent, we examined a series of aromatic aldehydes bearing a fluoro (**1b** and **1i**), a methoxy (**1h**), or a trifluoromethyl (**1e**) substituent. Similar EPR signals were observed for all substrates examined (Figure 2), indicating that ketyl radical formation is a common feature of these reactions. Variation of the substituents on the phenyl ring had little effect on the EPR lineshape, with one notable exception: when α-^13^C-labeled **1a** was used, significant line broadening arising from ^13^C hyperfine coupling was observed, consistent with the assignment of **III** as a TPP-bound, substrate-derived ketyl radical.

**Figure 2.**
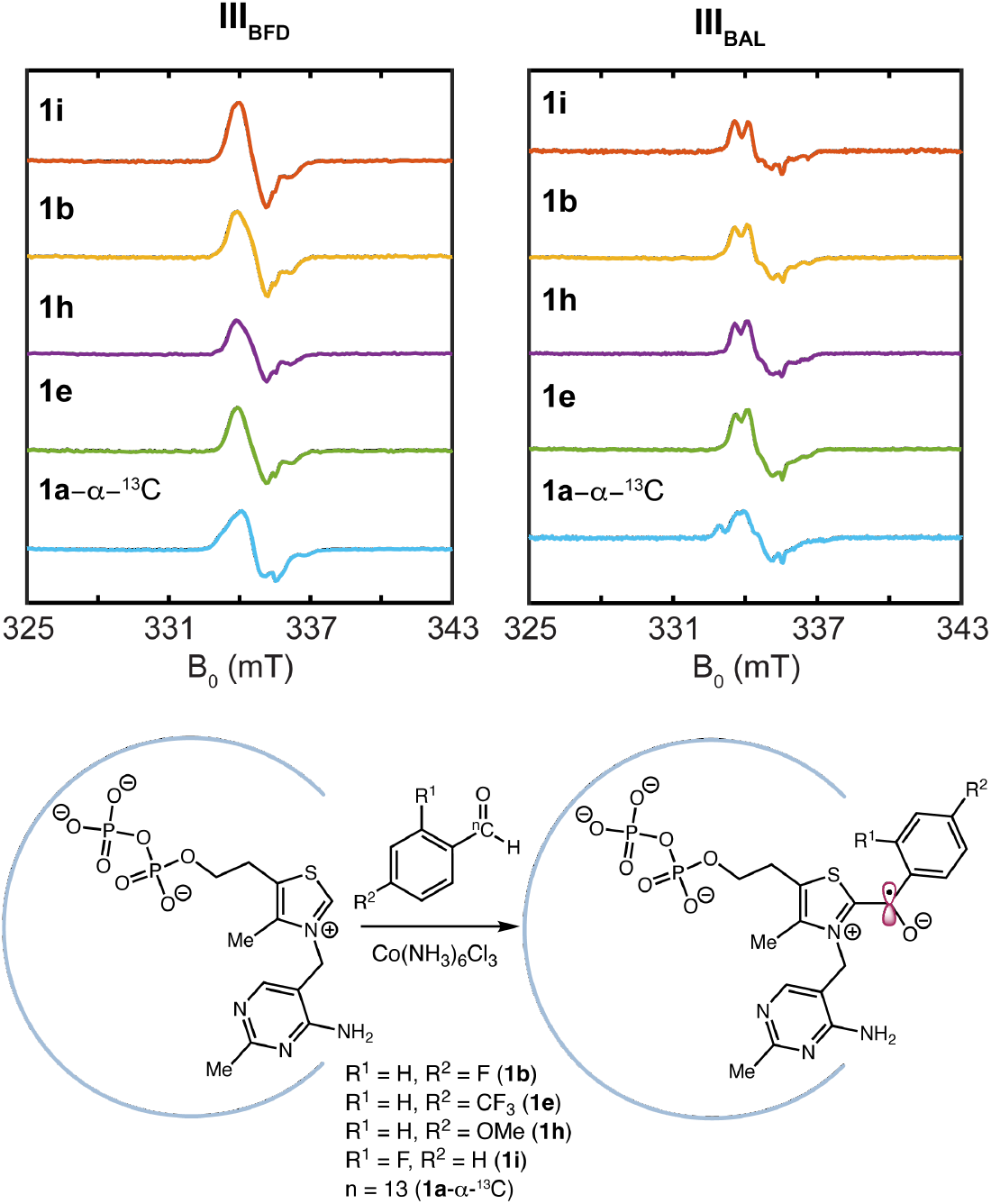
X-band EPR spectra of **III**_**BFD**_ (left) and **III**_**BAL**_ (right) generated from different aromatic aldehyde substrates.

To elucidate the origin of the distinct CW EPR lineshapes of **III**_**BFD**_ and **III**_**BAL**_, we further characterized these radicals using pulsed EPR and electron-nuclear double resonance (ENDOR) spectroscopy at higher microwave frequencies. At D-band (130 GHz), the *g* tensors of both radicals were fully resolved (Figure 3). Despite their distinct lineshapes observed at lower microwave frequencies, **III**_**BFD**_ and **III**_**BAL**_ exhibit very similar *g*-values. At Q-band (34 GHz), partial resolution of the *g* anisotropy, together with additional hyperfine splittings, was observed, particularly for **III**_**BAL**_. These splittings arise from a combination of ^1^H and ^14^N hyperfine interactions. The ^1^H couplings were confirmed by ENDOR spectroscopy (*vide infra*), whereas the ^14^N couplings give rise to additional splitting at Q-band and line broadening near *g*_3_ at D-band. The latter reflects a highly anisotropic hyperfine tensor associated with spin density in the 2p_z_ orbital of the thiazolium nitrogen. Simulations of the multifrequency EPR spectra provided excellent agreement with the experimental data for both radicals (Table 2). The simulated *g*-tensors, [2.0079, 2.0052, 2.0028] for **III**_**BFD**_ and [2.0085, 2.0056, 2.0029] for **III**_**BAL**_, exhibit *g*_3_ values close to the free-electron value (*g*_e_ = 2.0023), consistent with radicals possessing predominantly *p*_z_-type spin density. The larger shifts of *g*_1_ and *g*_2_, relative to typical carbon-centered radicals, likely arise from spin-orbit coupling contributions from the sulfur atom in the thiazolium ring. Overall, these experimental *g*-tensor values are in excellent agreement with the DFT-predicted values for TPP derived ketyl radicals (Table 2, *vide infra*).

**Table 2.** EPR Parameters for TPP-Dependent Enzyme-Bound Ketyl Radical Intermediates.

| entry | ketyl radical | g-tensor | $A\ ^1\text{H}^a$ | $A\ ^{31}\text{P}^a$ | $A\ ^{19}\text{F}^{a,b}$ | $A\ ^{14}\text{N}^{a,c}$ |
| --- | --- | --- | --- | --- | --- | --- |
| 1 | **III**<sub>BFD</sub> | [2.0079, 2.0052, 2.0028] | [7.6, 12.4, 7.1] ± 0.1;<br>$a_{\text{iso}} = 9.1$; $T = 1.7$ | [1.06, 0.67, 0.67] ± 0.01;<br>$a_{\text{iso}} = 0.80$; $T = 0.13$ | [2.5, 6.0, 2.5] ± 0.2<br>$a_{\text{iso}} = 3.7$; $T = 1.2$ | [∼, ∼, 24] ± 2 |
| 2 | **III**<sub>BFD</sub> DFT | [2.0073, 2.0042, 2.0000] | [9.2, 12.8, 6.2] |  | [6.3, 17.5, 6.3]<sup>d</sup> | [2, 2, 26.8] |
| 3 | **III**<sub>BAL</sub> | [2.0085, 2.0056, 2.0029] | [17.2, 21.3, 15.9] ± 0.1;<br>$a_{\text{iso}} = 18.1$; $T = 1.6$ | [1.00, 0.63, 0.65] ± 0.01;<br>$a_{\text{iso}} = 0.76$; $T = 0.12$ | [0.95, 1.45, 0.95] ± 0.05;<br>$a_{\text{iso}} = 1.1$; $T = 0.17$ | [∼, ∼, 31] ± 2 |
| 4 | **III**<sub>BAL</sub> DFT | [2.0086, 2.0050, 2.0018] | [17.9, 21.5, 14.7] |  | [1.7, 7.7, 2.5]<sup>d</sup> | [0, 0, 27.8] |
| 5 | TPP-COCH<sub>3</sub><sup>e</sup> | 2.008 |  | 0.51 |  |  |

<sup>a</sup>Hyperfine values in MHz; <sup>b</sup>4-CF<sub>3</sub>-C<sub>6</sub>H<sub>4</sub>CHO was used as the substrate; <sup>c</sup> $A_1$ and $A_2$ were too small to be accurately determined from simulations; <sup>d</sup>average value of three $^{19}\text{F}$ nuclei; <sup>e</sup>PFOR intermediate, see ref<sup>30a</sup>.

**Figure 3.**
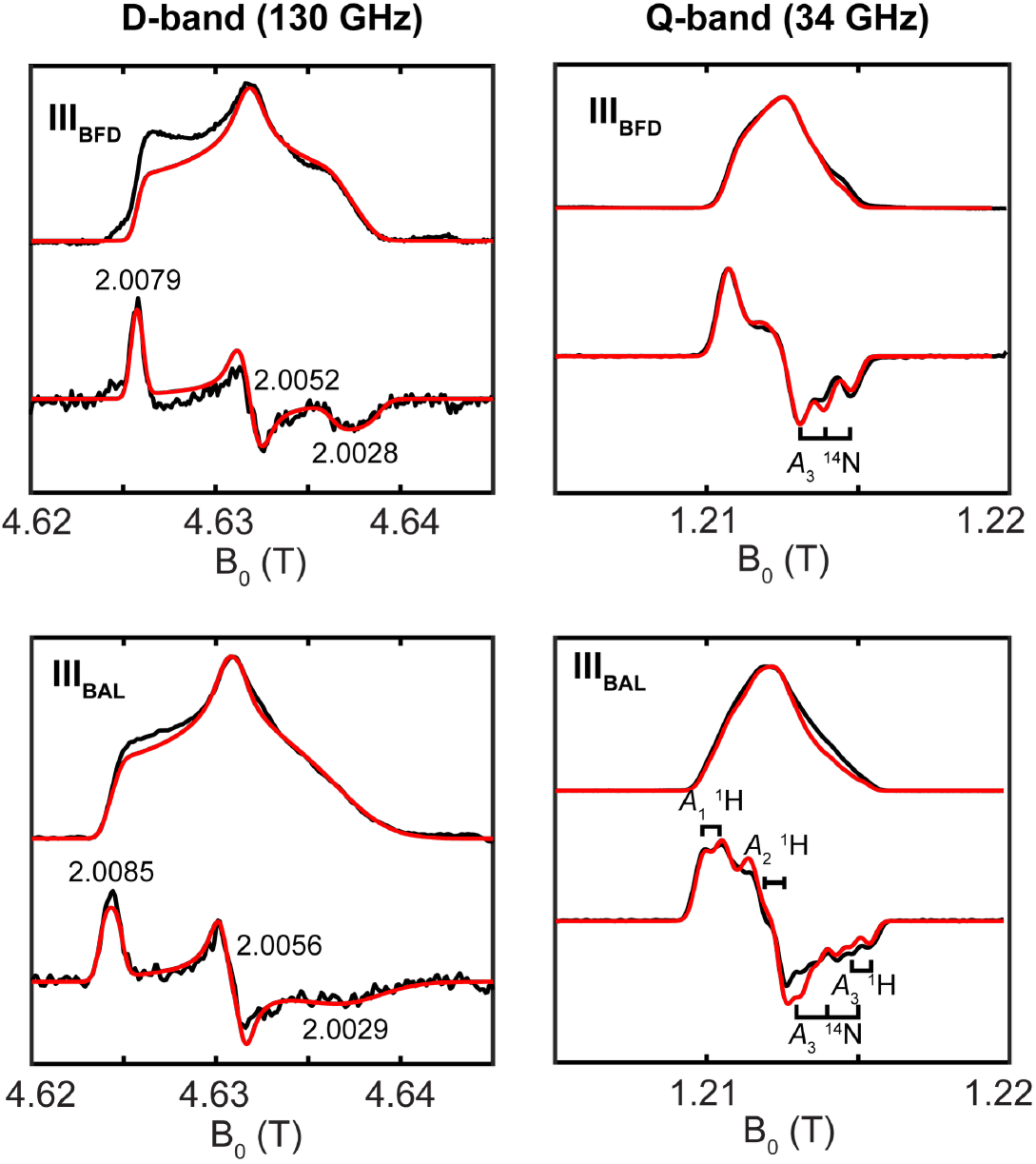
Electron spin echo detected field sweep and pseudomodulated EPR spectra (black traces) of **III**_**BFD**_ (top) and **III**_**BAL**_ (bottom) at D-band (left) and Q-band (right). Spectral simulations are in red traces. Spectrometer settings for Q-band: *T* = 55 K, τ = 300 ns, π/2 = 12 ns; and for D-band: *T* = 55 K, τ = 200 ns, π/2 = 40 ns. See Table 2 for simulation parameters.

Further mechanistic insight was obtained from ENDOR spectroscopic analysis. As shown in Figure 4A and Figure S1, Q-band Mims ENDOR spectra revealed small ^31^P hyperfine couplings of similar magnitude for both **III**_**BFD**_ and **III**_**BAL**_. Because the pyrophosphate group of TPP is the sole source of ^31^P in the sample, these hyperfine couplings provide direct spectroscopic evidence that the radical resides on a TPP-derived intermediate. Similar long-range ^31^P hyperfine couplings were previously reported for the TPP-derived ketyl radical intermediate in pyruvate:ferredoxin oxidoreductase (PFOR).^30a^

**Figure 4.**
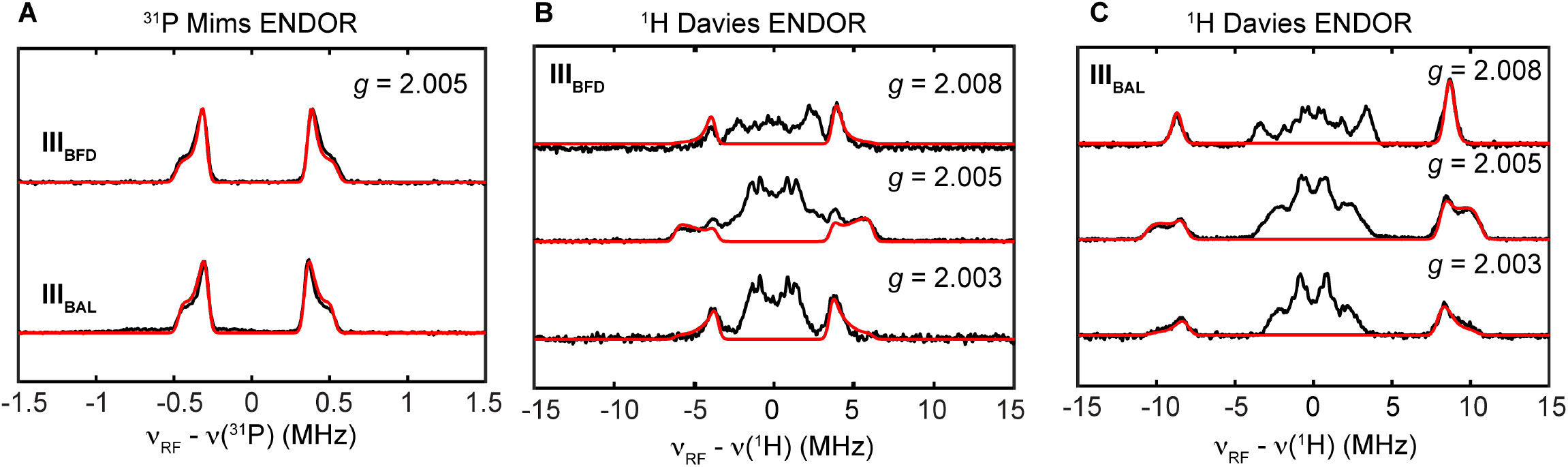
Q-band ENDOR spectra (black traces) and simulations (red traces). (A) Q-band ^31^P Mims ENDOR spectra of **III**_**BFD**_ and **III**_**BAL**_ collected at *g* = 2.005. See Figure S1 for field-dependent spectra. Field dependent ^1^H Davies ENDOR spectra of **III**_**BFD**_ (B) and **III**_**BAL**_ (C). ENDOR conditions: *T* = 55 K; inversion π pulse = 80 ns; τ = 300 ns, π/2 = 12 ns; RF pulse = 15 µs. See Table 2 for simulation parameters.

We subsequently performed field-dependent Q-band Davies ENDOR experiments to characterize the ^1^H hyperfine interactions (Figure 4). Strikingly, the ^1^H hyperfine couplings differed markedly between the two radicals. Spectral simulations gave ^1^H hyperfine tensors of [7.6, 12.4, 7.1] MHz (*a*_iso_ = 9.1 MHz) for **III**_**BFD**_ and [17.2, 21.3, 15.9] MHz (*a*_iso_ = 18.1 MHz) for **III**_**BAL**_, with the largest ^1^H coupling nearly twofold greater in **III**_**BAL**_. We propose that this substantial difference underlies the distinct X-and Q-band EPR lineshapes of **III**_**BFD**_ and **III**_**BAL**_, including the additional hyperfine splitting observed for **III**_**BAL**_.

The relatively isotropic ^1^H hyperfine couplings, together with the small dipolar components (*T* = 1.7 and 1.6 MHz for **III**_**BFD**_ and **III**_**BAL**_, respectively), suggest that these couplings arise primarily via a spin hyperconjugation mechanism from an sp^3^ β-proton adjacent to the delocalized spin center. Consistent with this assignment, performing the biocatalytic reactions in D_2_O-derived buffer had little effect on the ENDOR spectra of the resulting radicals (Figure 5A), indicating that the largest observed ^1^H hyperfine couplings do not originate from exchangeable protons and supporting the assignment of the ketyl oxygen in **III** as deprotonated. When perdeuterated **1a-***d*_5_ was used as the substrate, ^1^H ENDOR signals with smaller hyperfine couplings disappeared (Figure 5B), further confirming the identity of the radical as the TPP–**1a** adduct. However, the dominant ^1^H ENDOR signals persisted, excluding substrate phenyl protons as their origin. Together, these observations indicate that the largest ^1^H hyperfine interaction most likely arises from one of the methylene protons adjacent to the thiazolium ring. These results lead to an intriguing conclusion: although **III**_**BFD**_ and **III**_**BAL**_ correspond to the same TPP–**1a** ketyl radical consistent with the proposed mechanism, subtle differences in the active-site architectures of the two enzymes are sufficient to alter the radical geometry, thereby redistributing spin density and modulating the hyperfine interactions. Moreover, we note that the hyperfine couplings of the phenyl-derived ^1^H ENDOR signals lost upon phenyl deuteration show the opposite trend to that of the dominant ^1^H ENDOR signal. This reciprocal behavior suggests greater spin delocalization onto the phenyl ring in **III**_**BFD**_, accompanied by reduced spin density on the thiazolium ring, and *vice versa* for **III**_**BAL**_.

**Figure 5.**
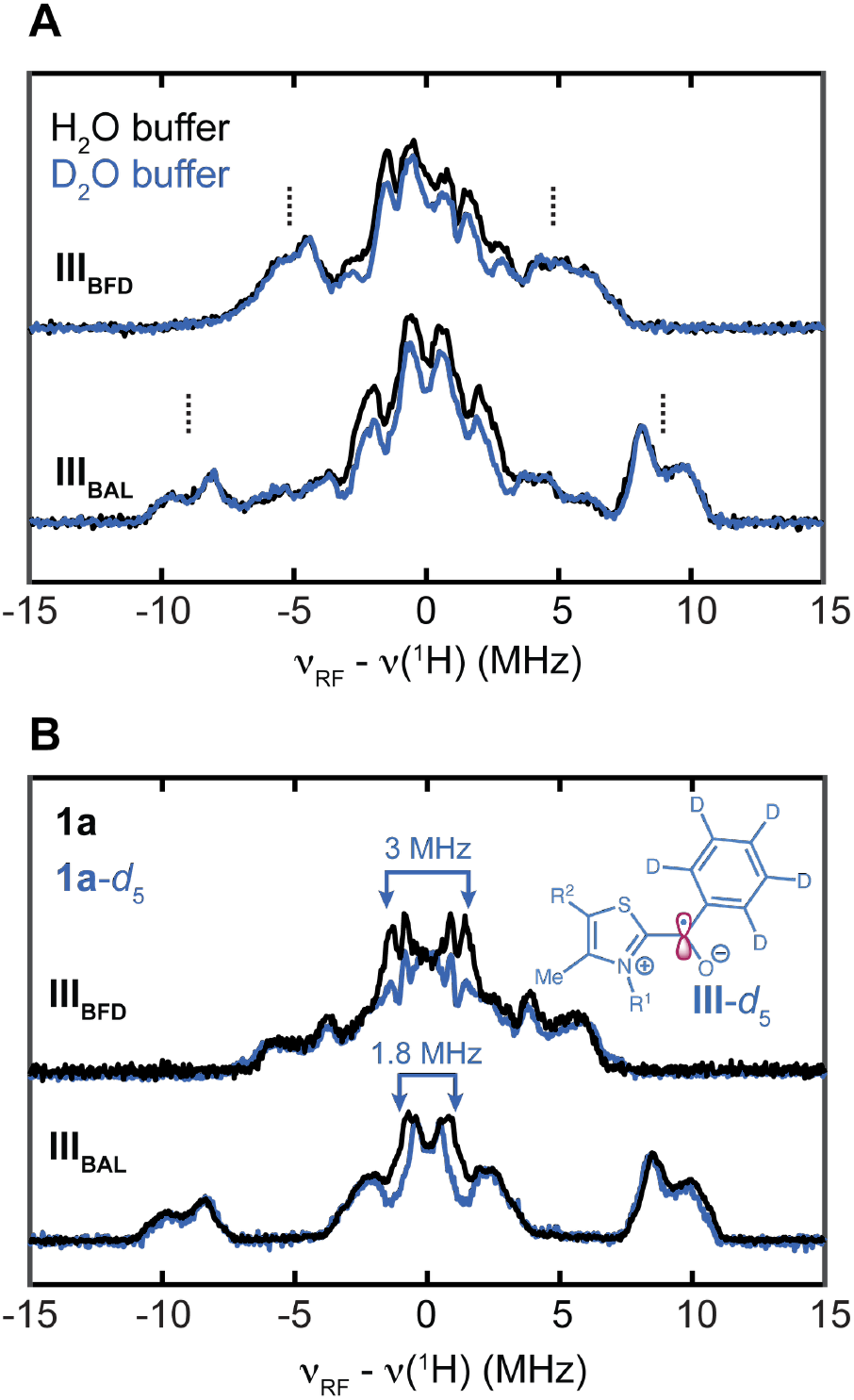
(A) Comparison of ^1^H Davies ENDOR spectra of **III**_**BFD**_ and **III**_**BAL**_ generated in H_2_O and D_2_O-based buffers. Major ^1^H ENDOR signals are indicated by dash lines. Here **1e** was used as the substrate. (B) Comparison of ^1^H Davies ENDOR spectra of **III**_**BFD**_ and **III**_**BAL**_ generated using **1a** and **1a**-*d*_5_ as substrates. Contributions from **1a** protons are indicated by arrows. ENDOR conditions are the same as in **Figure 4**.

To further probe spin delocalization onto the substrate-derived phenyl ring more directly, we employed 4-(trifluoromethyl)benzaldehyde (**1e**) as a spectroscopic probe. Because the three equivalent ^19^F nuclei are attached to the aromatic ring through an sp^3^ carbon (Figure 6A), they were expected to exhibit relatively isotropic hyperfine interactions that are readily detectable by ENDOR. At Q-band, however, the ^19^F ENDOR signals overlap extensively with the ^1^H signals because the nuclear gyromagnetic ratios of ^19^F and ^1^H are similar (B_0_ = 1.21 T; ν(^1^H) = 51.5 MHz; ν(^19^F) = 48.5 MHz). Nevertheless, the asymmetric Davies ENDOR lineshape observed for **III**_**BFD**_, but not **III**_**BAL**_, suggests contributions of unresolved ^19^F contributions (Figure S2).

**Figure 6.**
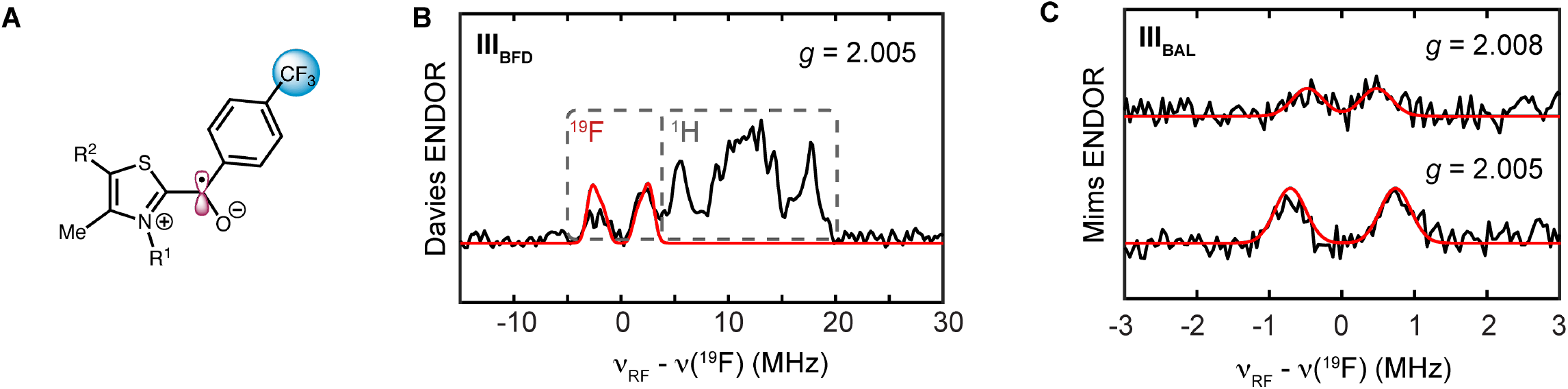
D-band ENDOR spectra. (A) Proposed structure of **III** formed using **1e** as the substrate. (B) D-band ^19^F Davies ENDOR of **III**_**BFD**_ collected at *g* = 2.005. (C) Field dependent ^19^F Mims ENDOR of **III**_**BAL**_ (C). ENDOR conditions: *T* = 55 K; inversion π pulse = 80 ns; τ = 200 ns, π/2 = 45 ns; RF pulse = 15 µs. See Table 2 for simulation parameters.

To resolve these interactions, we performed D-band Davies ENDOR experiments. As shown in Figure 6 and Figure S3, the higher magnetic field completely separated the ^19^F and ^1^H ENDOR signals (B_0_ = 4.63 T; ν (^1^H) = 197 MHz; ν (^19^F) = 185 MHz). Spectral simulations provided a ^19^F hyperfine tensor of [2.5, 6.0, 2.5] MHz (*a*_iso_ = 3.7 MHz; *T* = 1.2 MHz) for **III**_**BFD**_. In contrast, the corresponding ^19^F hyperfine coupling in **III**_**BAL**_ were substantially smaller and could only be resolved by D-band Mims ENDOR, giving [0.95, 1.45, 0.95] MHz (*a*_iso_ = 1.1 MHz; *T* = 0.17 MHz). The approximately threefold reduction in ^19^F hyperfine coupling indicates significantly less spin delocalization onto the phenyl ring in **III**_**BFD**_ than in **III**_**BAL**_, providing direct evidence that the enzyme environment subtly modulates the electronic structure of an otherwise identical ketyl radical intermediate.

To better understand how the enzyme environment modulates the electronic structure of the ketyl radical intermediate, we performed DFT calculations to evaluate the hyperfine coupling constants for the ketyl radical derived from TPP and 4-(trifluoromethyl)benzaldehyde (**1e**) over a range of conformations. Specifically, as shown in Figure 7, the N–C1–C2–O dihedral angle was systematically varied, while the TPP moiety was held fixed to mimic the constrained cofactor geometry within the enzyme active site. This conformational search identified two distinct structures that reproduce the experimentally determined largest ^1^H hyperfine coupling of **III**_**BFD**_ and **III**_**BAL**_, respectively. In both conformations, the largest ^1^H hyperfine couplings originate from the methylene protons adjacent to the thiazolium nitrogen, consistent with substantial spin density localized at this position and the spin hyperconjugation mechanism proposed above. Although accurate prediction of ^19^F hyperfine couplings is inherently more challenging because the isotropic (Fermi contact) contribution is highly sensitive to the spin density at the fluorine nucleus, the calculated spin density distribution correctly reproduces the experimentally observed trend in ^19^F hyperfine couplings between **III**_**BFD**_ and **III**_**BAL**_. Importantly, the two DFT-predicted conformations fit well within the crystallographic activesite pocket of BFD and BAL (Figure S4), indicating their relevance to radical biocatalysis. In BFD, H281 points toward the aromatic ring of the aldehyde substrate, whereas in BAL, T481 lies in proximity to the thiazolium ring. Mutations at these sites may preferentially favor the respective radical conformation shown in Figure 7. Together, these results demonstrate that the enzyme-dependent conformational constraints imposed on the TPP-bound ketyl radical redistribute spin density, providing a molecular basis for the distinct hyperfine couplings observed by EPR spectroscopy.

**Figure 7.**
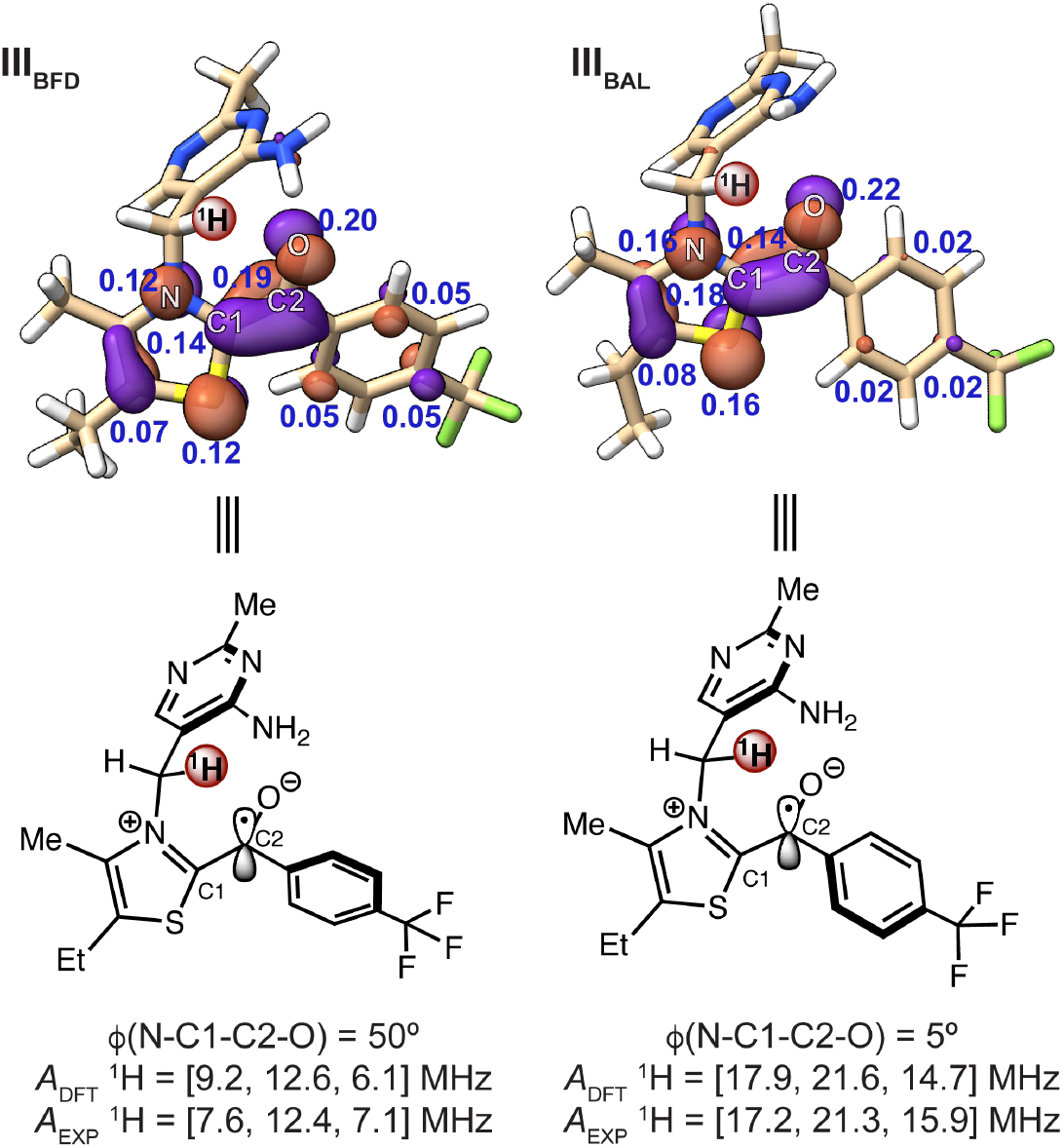
Structure models of **III**_**BFD**_ (left) and **III**_**BAL**_ (right). Calculated Löwdin spin densities are shown in blue. Isosurfaces indicate a contour value of 0.05. The protons with largest ^1^H hyperfine coupling are labeled in red spheres. See Table 2 for the summary of DFT-predicted EPR parameters.

The multifrequency EPR and ENDOR results presented here provide a unique opportunity to correlate the electronic structures of the ketyl radical intermediates with the distinct catalytic behaviors of BFD and BAL in radical-mediated C–C coupling reactions. Although **III**_**BFD**_ and **III**_**BAL**_ correspond to the same TPP–substrate ketyl radical, their spectroscopic signatures reveal measurable differences in spin distribution arising from their distinct active-site environments.

The most direct evidence comes from the ^1^H and ^19^F hyperfine couplings. **III**_**BAL**_ exhibits substantially larger ^1^H hyperfine couplings associated with the thiazolium methylene group, indicating increased spin density on the TPP cofactor. In contrast, the approximately threefold larger ^19^F hyperfine coupling observed for **III**_**BFD**_ demonstrates significantly greater spin delocalization into the substrate-derived phenyl ring. Together, these observations indicate that the unpaired electron is distributed differently in the two enzymes despite their nearly identical chemical compositions. The small differences in the *g* tensors further support subtle changes in electronic structure, rather than formation of different radical intermediates. Consistent with these spectroscopic observations, DFT calculations identified distinct ketyl radical conformations that reproduce the experimental hyperfine coupling patterns of **III**_**BFD**_ and **III**_**BAL**_, demonstrating that enzyme-specific conformational constraints modulate the electronic structure of the radical intermediate.

These electronic differences provide a plausible explanation for the contrasting catalytic activities of BFD and BAL. In BFD, greater spin delocalization toward the substrate implies increased spin density at the benzylic ketyl carbon, the site that subsequently undergoes intermolecular radical coupling. As a consequence, **III**_**BFD**_ possesses a more substrate-centered singly occupied molecular orbital (SOMO), facilitating SOMO–SOMO overlap with the incoming benzylic radical during C–C bond formation. Conversely, in BAL, a larger fraction of the unpaired spin remains localized on the thiazolium ring of the TPP cofactor. This cofactor-centered spin localization stabilizes the ketyl radical while diminishing radical character on the substrate fragment, thereby reducing its propensity for intermolecular radical C–C coupling. Thus, enzyme-dependent redistribution of spin density provides a mechanistic basis for the divergent catalytic outcomes of these closely related TPP-dependent enzymes.

Our findings establish a correlation between the electronic structure of a common ketyl radical intermediate and enzyme function, demonstrating that spin-density distribution serves as an electronic descriptor of catalytic competence in TPP-dependent radical biocatalysis. More broadly, these results reveal that protein scaffolds can regulate radical reactivity not by changing the chemical identity of the intermediate, but by subtly tuning the electronic structure through active-site interactions. This mechanistic insight suggests that engineering the electronic differences of protein-bound radical intermediates represents a potentially general strategy for controlling enzyme reactivity and selectivity in radical biocatalysis. Interestingly, the radical SAM enzymes BzaF and ThiC also share the same initial radical intermediate derived from their common substrate, 5-aminoimidazole ribotide. The geometric and electronic structure of this radical could also be differentially tuned by the respective enzyme active sites, directing its subsequent reactivity along distinct pathways and ultimately leading to the completely different products generated by BzaF and ThiC.^32, 33^

Finally, with the optimized biocatalytic reaction conditions, we surveyed the scope of the oxidative radical C–C coupling catalyzed by *Pp*BFD variants (Table 3). It was found that benzaldehydes bearing halogen substituents, including a *para*-fluoro (**4b**), a *para*-chloro (**4c**), and a *para*-bromo group (**4d**), were well tolerated, affording the corresponding ketone products with excellent enantioselectivity. Substrates containing electron-withdrawing groups, such as a *para*-trifluoromethyl (**4e**) and a *para*-trifluoromethoxy (**4f**) group, as well as electron-donating groups, including a *para*-methyl (**4g**) and a *para*-methoxy group (**4h**), were also readily accommodated. Additionally, benzaldehyde possessing an *ortho*-fluorine (**4i**) substituent could also be transformed with good enantioselectivity, although with a modest yield. Furthermore, a range of secondary alkyltrifluoroborate salts were readily transformed. Aryl-substituted trifluoroborates bearing a fluorine (**4j**), a chlorine (**4k**) and a bromine (**4l**) substituent underwent efficient oxidative coupling. Organotrifluoroborate substrates bearing a trifluoromethyl group (**4m**) and a methyl group (**4n** and **4o**) were also transformed with excellent enantioselectivity. Moreover, secondary alkylboron reagents with an extended aliphatic substituent, including an indanyl (**4p**) and ethyl (**4q**) and *n*-propyl (**4r**) substituents, readily underwent this oxidative photobiocatalytic coupling. Collectively, these results further demonstrated the synthetic utility of engineered BFD variants for enabling oxidative radical C–C bond formation inaccessible to TPP-dependent BAL enzymes.

**Table 3.**
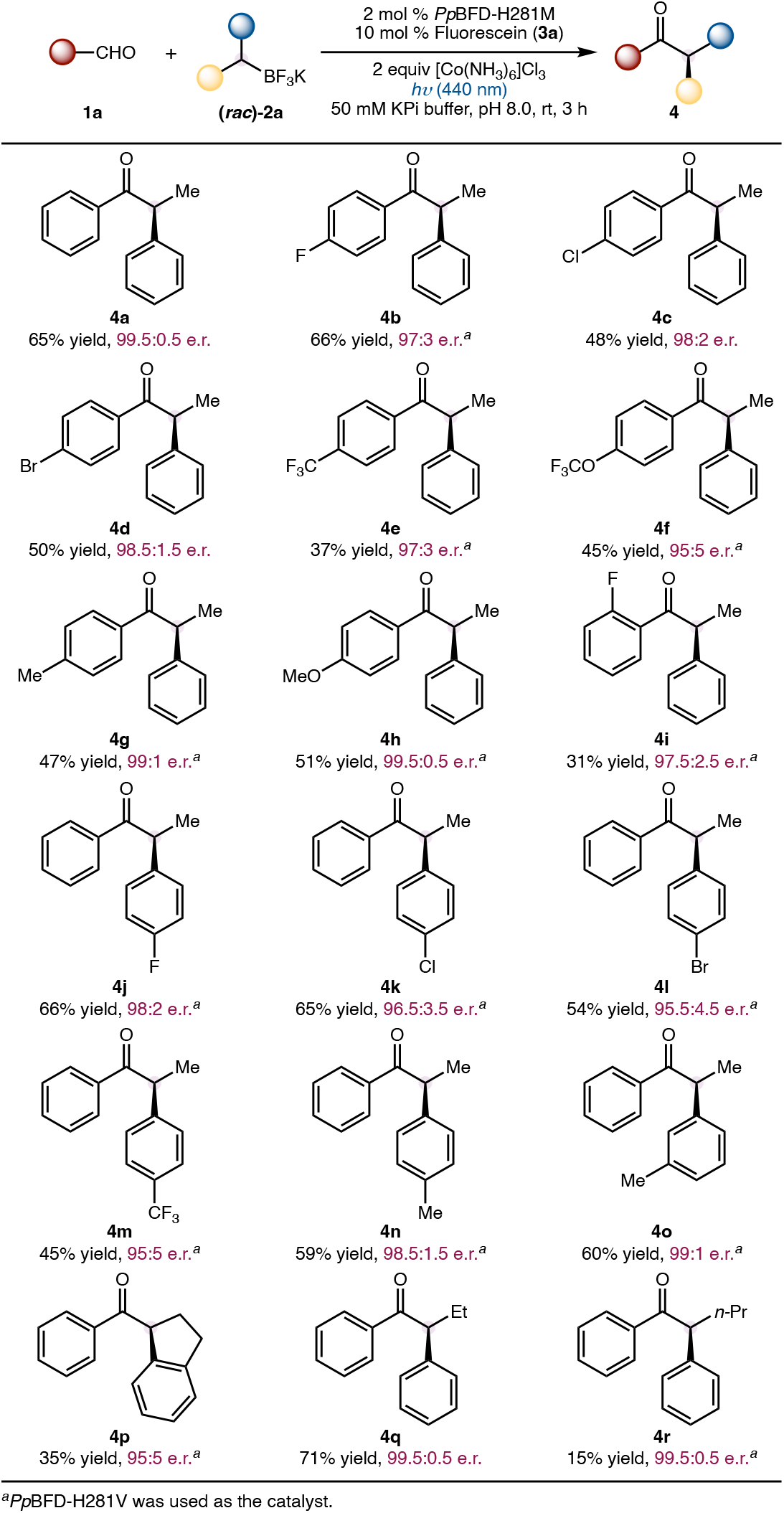
Substrate Scope of TPP Enzyme-Catalyzed Asymmetric Oxidative Radical C–C Coupling.

## CONCLUSIONS

In summary, we have developed an oxidative photobiocatalytic asymmetric radical C–C coupling catalyzed by engineered TPP-dependent BFD variants and used this reaction to investigate the nature of the enzyme-bound ketyl radical intermediates that underpin TPP-dependent radical biocatalysis. Freeze-quench multifrequency EPR and ENDOR spectroscopy directly identified the catalytically generated TPP-bound ketyl radical and provided the detailed spectroscopic characterization of this central intermediate in non-native TPP radical enzymology. Although BFD and BAL generate the same ketyl radical intermediate, their EPR signatures reveal distinct electronic structures arising from differences in the enzyme active-site environment. Combined spectroscopic and DFT analyses demonstrate that enzyme-dependent conformational constraints redistribute spin density between the TPP cofactor and the substrate fragment, providing a molecular basis for the divergent radical reactivities of these closely related TPP-dependent enzymes. Greater substrate-centered spin delocalization in BFD correlates with enhanced intermolecular radical C–C coupling, whereas increased cofactor-centered spin localization in BAL is associated with diminished coupling reactivity. Beyond elucidating the mechanism of TPP-dependent radical biocatalysis, this work establishes a broader principle for enzyme catalysis: protein scaffolds can regulate radical reactivity by tuning the electronic structure of a common radical intermediate rather than by changing its chemical identity. The ability to experimentally resolve these subtle electronic differences by multifrequency EPR and ENDOR spectroscopy provides a new framework for understanding enzyme-controlled radical chemistry, demonstrating spin-density distribution engineering as a promising strategy for developing new radical biocatalysts with improved reactivity and selectivity.

## ASSOCIATED CONTENT

### Supporting Information

The Supporting Information is available free of charge on the ACS Publications website.

Additional experimental and computational results, EPR spectra, characterization data, HPLC traces and NMR spectra (PDF)

## AUTHOR INFORMATION

### Notes

The authors declare no competing financial interest.

## ACKNOWLEDGMENT

We are grateful to the Army Research Office (W911NF-23-1-0182 to Y.Y.) and the National Institutes of Health (R35GM126961 to R.D.B.) for financial support. Y.Y. is an Alfred P. Sloan Research Fellow (FG-2024-22244), a Camille Dreyfus Teacher-Scholar Awardee (TC-25-084), a David & Lucile Packard Fellow (2023-76169) and a Howard Hughes Medical Institute Freeman Hrabowski Scholar. Computations were carried out at the Data and the Advanced Cyberinfrastructure Coordination Ecosystem: Services & Support (ACCESS) Program, supported by NSF award CHE-260005.

